# CETSA^®^ MS profiling for a comparative assessment of FDA approved antivirals repurposed for COVID-19 therapy identifies Trip13 as a Remdesivir off-target

**DOI:** 10.1101/2020.07.19.210492

**Authors:** Tomas Friman, Alexey Chernobrovkin, Daniel Martinez Molina, Laurence Arnold

## Abstract

The reuse of pre-existing small molecules for a novel emerging disease threat is a rapid measure to discover unknown applications for previously validated therapies. A pertinent and recent example where such strategy could be employed is in the fight against COVID-19. Therapies designed or discovered to target viral proteins also have off-target effects on the host proteome when employed in a complex physiological environment. This study aims to assess these host cell targets for a panel of FDA approved antiviral compounds including Remdesivir, using the cellular thermal shift assay (CETSA^®^) coupled to mass spectrometry (CETSA MS) in non-infected cells. CETSA MS is a powerful method to delineate direct and indirect interactions between small molecules and protein targets in intact cells. Biologically active compounds can induce changes in thermal stability, in their primary binding partners as well as in proteins that in turn interact with the direct targets. Such engagement of host targets by antiviral drugs may contribute to the clinical effect against the virus but can also constitute a liability. We present here a comparative study of CETSA molecular target engagement fingerprints of antiviral drugs to better understand the link between off-targets and efficacy.

## Introduction

The COVID-19 pandemic has seen a significant worldwide effort to reuse or repurpose preexisting therapies in order to combat the emerging viral threat. There have been numerous studies reported using a variety of technologies in efforts to screen panels of pre-validated molecules, many repurposed from viral therapies^1–5^. These studies are conducted in the hope that efficacy against the SARS-CoV-2 virus may be discovered, whilst avoiding the lengthy yet essential drug discovery pipeline that even with modern standards typically takes several years from hit and target identification, to reach clinical testing of a lead candidate drug molecule^6^.

Utilising a preexisting molecule has a significantly lower risk than rapidly developing novel chemistry, since it has already successfully navigated the prerequisite safety and toxicologic testing for use in humans. However, the original purpose of the small molecule may have undescribed off-target effects that are deemed to be tolerable when weighed against therapeutic benefit. These effects, potentially caused by drug-protein interactions, are often poorly understood^7^.

For example, many antiviral compounds are structural analogues of nucleoside triphosphates (NTPs) that have diverse biological properties and therapeutic consequences since nucleotides have an essential role in virtually all biological processes^8^. Therefore, given the abundance of nucleotide interacting proteins in the host cell, off-target interacting proteins, or an imbalance of the cellular nucleotide pool would be an expected consequence of utilising nucleotide analogues in therapy^9^.

The persistent and fundamental problem of host off-target effects arise from using a molecule to disrupt viral biology, whilst simultaneously exposing the host biology to the same chemical challenge. Methods to describe the severity of hitting off targets, rely upon in vitro and in vivo assessment or the presentation of a phenotype that can be assessed as to whether acceptable or not. But this requires knowledge and the ability to measure non-intended target biology. For example, Remdesivir is known to have an efficacy of 100nM for the viral polymerase its intended target, and 500-fold less efficacious against human polymerases^10^. It has previously not been established which other proteins may interact with and nor whether these potential interactions would elicit a response with a measurable output using conventional means.

In light of this, traditional off-target investigation relies on known functions or activities which as a prerequisite require the host proteins responsible for these activities to be studied in bias. Methods that are independent of activity and in an unbiased way report on compound interaction against the entire proteome, have only in recent years been established^11,12^. The CEllular Thermal Shift Assay (CETSA) is a powerful technique to detect protein ligand interactions in cells^13^. Coupled with mass spectrometry (MS) as a readout, CETSA MS is a technique employed in the identification of off-target effects in proteome-wide studies observing the thermal stabilization or destabilization of endogenous proteins and downstream effects after matrix and compound incubation. The method is being increasingly employed in both mechanism of action (MoA) studies and to identify primary and off-targets of candidate drug molecules. For example, quinine and drug target interactions in *Plasmodium falciparum* ^14,15^. In this study, we screened a panel of drugs using the CETSA MS format on HepG2 cells to identify host proteins as hopeful starting points for further research and possible inroads into the improvement or development of fortuitous therapies for SARS-CoV-2 infection.

Given the intense global interest in searching for a viable therapy combined with the wide accessibility to information sources and even raw data, efforts from a wide variety of groups have been well documented in both the scientific and non-scientific media. The inclusion of compounds for this study was directed around prominent molecules discussed in the literature and adopted for clinical trials in the earlier phases of the worldwide pandemic, namely Remdesivir and Hydroxychloroquine. The study was bolstered by the edition of other compounds repurposed from a variety of anti-viral classes including retroviral reverse transcriptase and protease inhibitors that were available for expeditious purchase from commercial sources^16–19^.

This study investigates compound effects on uninfected whole HepG2 cells. Understanding how the molecule reacts in an environment containing both viral and host cell proteins is not beyond the technique, but outside of the capacity and scope for this study that was completed utilising a preexisting in vitro platform with a per compound acquisition time of approximately ~6 hours.

## Materials and Methods

### Cell culture

The human cell line HepG2 was procured from ATCC and cultured until 70% confluency in collagen-coated flasks. The cells were cultured in DMEM/F12 (without phenol red) (ThermoFisher) supplemented with 10% FBS (ThermoFisher), 5 mM sodium pyruvate (ThermoFisher), 1X NEAA (ThermoFisher) and PEST (ThermoFisher). Cells were detached using 5 ml of Tryp-LE (Thermo Fisher Scientific), pelleted, washed with Hank’s Balanced Salt solution (HBSS, Thermo Fisher Scientific), and pelleted again. Cell viability was measured with trypan blue exclusion and cells with a viability above 90% were used for these experiments. Cell pellets were resuspended in medium-free incubation buffer (20 mM HEPES, 138 mM NaCl, 5 mM KCl, 2 mM CaCl_2_, 1 mM MgCl_2_, pH 7.4) for further use as a 2x cell suspension.

### Compound handling

All compounds were acquired from commercial sources as powder stocks and reconstituted in Dimethyl sulphoxide (DMSO) or aqueous buffer dependent on manufacturer recommended solubility values. DMSO concentration was normalized across all samples to a final concentration of 1% v/v.

### Compressed CETSA MS experiment

The cell suspension was divided into 32 aliquots (22 test compounds, 2x methotrexate (1^st^ positive control), 2x vincristine (2^nd^ positive control) and 4x negative/vehicle controls) in Eppendorf tubes and mixed with an equal volume of either of the test compound or controls at 2x final concentration in the experimental buffer. The resulting final concentration of the compounds was 30 μM; 1% DMSO was used as a vehicle control. Incubations were performed for 60 minutes at 37°C with end-over-end rotation. Viability after 1 hour of incubation with each compound were greater than 90%.

Each of the treated cell suspensions was further divided into 12 aliquots that were all subjected to a heat challenge for 3 minutes, each at a different temperature between 44 and 66°C. After heating, all temperature points for each test condition were pooled to generate 32 individual (compressed) samples.

Precipitated proteins were pelleted by centrifugation at 30 000 x g for 20 minutes and supernatants constituting the soluble protein fraction were kept for further analysis.

The experiment was performed over three independent biological replicates.

### Protein digestion

The total protein concentration of the soluble fractions were measured by Lowry DC assay (BioRad). From each soluble fraction, a volume containing an equivalent of 20μg of total protein was taken for further sample preparation.

Samples were subjected to reduction and denaturation with tris(2-carboxyethyl)phosphine (TCEP) (Bond-breaker, Thermo Scientific) and RapiGest SF (Waters), followed by alkylation with chloroacetamide. Proteins were digested with Lys-C (Wako Chemicals) and trypsin (Trypsin Gold, Promega).

### TMT-labeling of peptides

After complete digestion had been confirmed by nanoLC-MS/MS, samples were labelled with 16-plex Tandem Mass Tag reagents (TMTpro, Thermo Scientific) according to the manufacturer’s protocol.

Labeling reactions were quenched by addition of a primary amine buffer and the test concentrations and room temperature control samples were combined into TMT16-plex sets such that each TMT16-multiplex set contained 12 test compounds, two positive control samples (MTX+Vincristine) and two negative controls (1% DMSO). The labelled samples were subsequently acidified and desalted using polymeric reversed phase chromatography (Oasis, Waters). LC-MS grade liquids and low-protein binding tubes were used throughout the purification. Samples were dried using a centrifugal evaporator.

### LC-MS/MS analysis

For each TMT16-multiplex set, the dried labelled sample was dissolved in 20 mM ammonium hydroxide (pH 10.8) and subjected to reversed-phase high pH fractionation using an Agilent 1260 Bioinert HPLC system (Agilent Technologies) over a 1.5 x 150 mm C18 column (XBridge Peptide BEH C18, 300 Å, 3.5 μm particle size, Waters Corporation, Milford, USA). Peptide elution was monitored by UV absorbance at 215 nm and fractions were collected every 30 seconds into polypropylene plates. The 60 fractions covering the peptide elution range were evaporated to dryness, ready for LC-MS/MS analysis.

From the fractions collected, 30 pooled fractions were analyzed by high resolution nano LC-MS/MS on Q-Exactive HF-X Orbitrap mass spectrometers (Thermo Scientific) coupled to high performance nano-LC systems (Ultimate 3000 RSLC Nano, Thermo Scientific).

MS/MS data was collected using higher energy collisional dissociation (HCD) and full MS data was collected using a resolution of 120 K with an AGC target of 3e6 over the m/z range 375 to 1500. The top 15 most abundant precursors were isolated using a 1.4 Da isolation window and fragmented at normalized collision energy values of 35. The MS/MS spectra (45 K resolution) were allowed a maximal injection time of 120 ms with an AGC target of 1e5 to avoid coalescence. Dynamic exclusion duration was 30 s.

### Protein identification and quantification

Protein identification was performed by database search against 95 607 human protein sequences in Uniprot (UP000005640, download date: 2019-10-21) using the Sequest HT algorithm as implemented in the ProteomeDiscoverer 2.4 software package. Data was re-calibrated using the recalibration function in PD2.4 and final search tolerance settings included a mass accuracy of 10 ppm and 50 mDa for precursor and fragment ions, respectively. A maximum of 2 missed cleavage sites were allowed using fully tryptic cleavage enzyme specificity (K, R, no P). Dynamic modifications were; oxidation of Met, and deamidation of Asn and Gln. Dynamic modification of protein N-termini by acetylation was also allowed. Carbamidomethylation of Cys, TMTpro-modification of Lysine and peptide N-termini were set as static modifications. For protein identification, validation was done at the peptide-spectrum-match (PSM) level using the following acceptance criteria; 1 % FDR determined by Percolator scoring based on Q-value, rank 1 peptides only. For quantification, a maximum co-isolation of 50 % was allowed. Reporter ion integration was done at 20 ppm tolerance and the integration result was verified by manual inspection to ensure the tolerance setting was applicable. For individual spectra, an average reporter ion signal-to-noise of >20 was required. Only unique or razor peptides were used for protein quantification.

### Data analysis

Quantitative results were exported from Proteome Discoverer as tab-separated files and analyzed using R version 4.0.2 software. Protein intensities in each TMT channel were log2-transformed and normalized by subtracting median value per each TMT sample and each TMT channel (column-wise normalization). For each protein and each compound, thermal stability changes were assessed by comparing normalized log2-transformed intensities to DMSO treated control using moderated t-test implemented in “limma” R-package version 3.44.1^20^.

## Results

We have applied the cellular thermal shift assay combined with quantitative LC-MS based proteomics (CETSA MS) to profile compound-induced protein thermal stability changes for 22 compounds in intact HepG2 cells. The experiments were performed in compressed, or one-pot format^21^. HepG2 cells were treated with 30μM of each compound and incubated for 60 minutes at 37°C in serum-free salt-based medium. After compound treatment, cell suspensions were divided into 12 aliquots followed by heat shock treatment at 12 temperatures (44-66°°C) for 3 min. After heat treatment, samples were pooled and aggregated proteins were removed by centrifugation. Resulting protein abundance in the soluble fraction corresponded to the area under the protein’s melting curve. The experiment was repeated to yield three biological replicates. Single compound concentration and application of compressed (one-pot) experimental design allowed for reliable protein stability changes assessment at relatively high throughput of ~6 hours acquisition time per compound. The studied compounds will be incorporated in a larger (>200 compounds) initiative to establish CETSA based molecular fingerprints of a diverse set of compounds.

**Figure 1.**
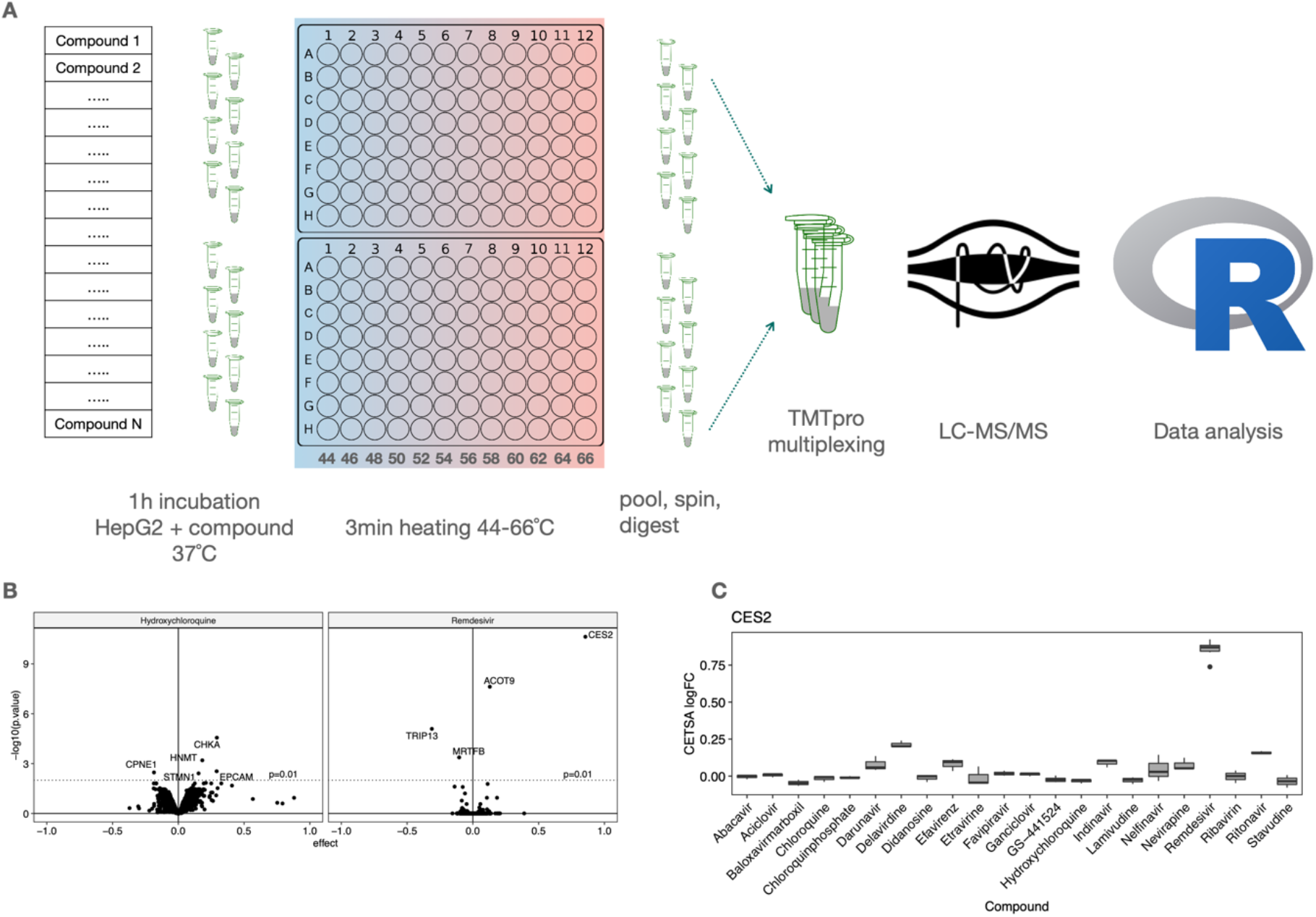
(A) Design of the experiment for CETSA MS profiling of 22 compounds in intact HepG2 cells. (B) Volcano plots summarising proteins found to be stabilized/destabilized upon treatment of HepG2 cells with Hydroxychloroquine (left) and Remdesivir (right). (C) Boxplot representation showing stability changes of cocaine esterase CES2 relative to the vehicle control for all 22 compounds analysed.

Proteins were quantified via isobaric labelling liquid chromatography coupled to mass-spectrometry (LC-MS). The resulting dataset covers more than 8,000 protein groups, of them 5,873 protein groups were reliably quantified in more than 17 out of 22 treatments with at least two unique peptides.

In order to assess compound induced protein thermal stability changes, for each treatment we compared log2-transformed and normalized intensities to the corresponding vehicle controls. For all 22 compounds tested only 34 proteins were found to be significantly changed (stabilized or destabilized) upon treatment with at least one compound (Figure 2).

Remdesivir, Ritonavir, Baloxavir marboxil and Chloroquines demonstrate distinct proteome responses and form individual clusters in hierarchical clustering. The remaining compounds are represented in Cluster 2.

**Figure 2.**
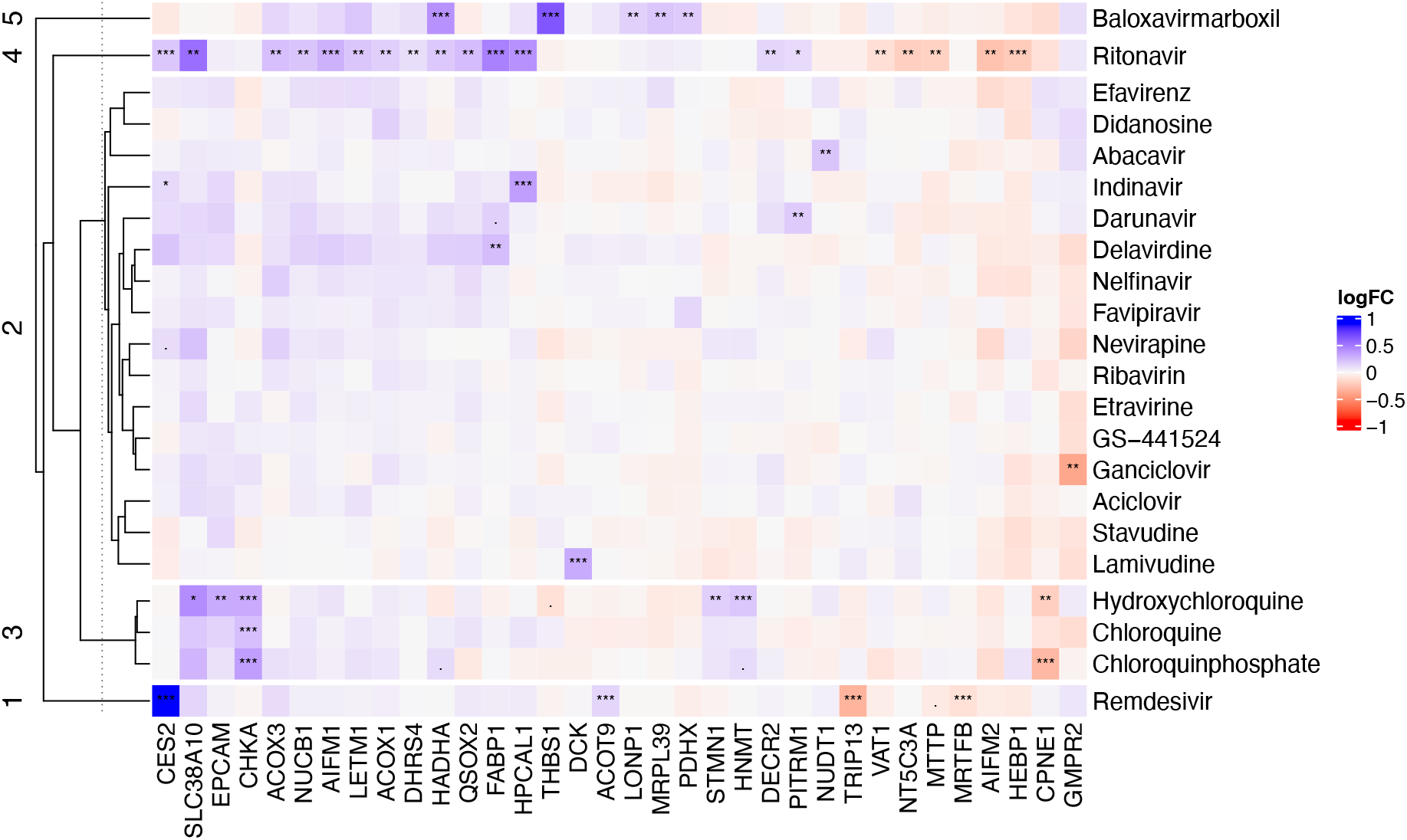
Heatmap of compound induced protein thermal stability changes in HepG2 cells treated with different antiviral compounds. Proteins found to be significantly changed (p≤0.01) in at least one compound included into the plot.

Remdesivir, being one of several nucleoside analogues in our panel, shows a clear hit for Carboxylesterase 2 (CES2) which could be involved in the metabolism of the molecule.

Although hydrolysis of the ester is reportedly by cathepsin A and carboxylesterase 1 (CES1)^22^, CES2 has high abundance in liver tissue. Also, Acyl-coenzyme thioesterase 9 (ACOT9) as well as Diphthine methyl ester synthase (DPH5) - albeit less obvious, showed stability shift with treatment, both of these proteins are known to bind esters, similar to the activity of CES2. Given during the activation of the prodrug includes an intracellular esterase hydrolysis step, an interaction is not surprising.

In contrast, and most notable from this study is the destabilization of Pachytene checkpoint protein 2 homolog (TRIP13). Trip13 is a hexameric AAA+ ATPase and a key regulator in chromosome recombination and structural regulation, such as crossing over and DNA double strand breaks^23^. Trip13 is essential in the spindle assembly checkpoint and is expressed in a number of human cancers where its reduction has been linked with effects on proliferation and hence therapeutic benefit^24^. It is plausible that Remdesivir, in its fully synthesized triphosphate form is competitive with endogenous ATP binding with Trip13, disrupting or affecting multimerization with itself or downstream on the spindle assembly complex.

Interestingly, GS-441524, a metabolite of Remdesivir had no significant hits in this study. There could be multiple explanations for this, but in this case, it is established that unfavorable compound properties of GS-441524 result in limited cellular uptake. Especially when a 60-minute incubation protocol is considered. In our experience, addition of nucleosides often has impact on several proteins involved in cellular nucleoside homeostasis.

As apparent from Figure 2, the Chloroquines (Hydroxychloroquine, Chloroquine and Chloroquine Phosphate) comprise their own cluster. Despite having clear function on the endosomal processes, the hits identified for hydroxychloroquine do not appear to follow an obvious pathway response e.g. vesicle proteins, vesicle lumen proteins (including ER Golgi), extracellular proteins, ion channels and transporters. It is possible that the identified hits are best representatives for the technique from their respective pathways. However, there are no known literature sources linking these targets to chloroquine and hydroxy version to activity found.

Despite this, a common hit between all three chloroquine derivatives tested was Choline Kinase alpha (CHKA) that has a key role in phospholipid biosynthesis. Another common hit between the hydroxy and Chloroquine phosphate forms is Copine 1 (CPNE1) a calcium dependent phospholipid binding protein that plays a role in calcium-mediated intracellular processes^25^. Other significant hits are <u>Histamine N-methyltransferase (HNMT)</u>, Epithelial cell adhesion molecule (EPCAM), and Stathmin (STMN1).

As concluded earlier, both Remdesivir and the Chloroquines stand out as separate clusters with no other antiviral compounds having similar response patterns. However, Ritonavir, a protease inhibitor and Baloxavir marboxil also stand out with unique response patterns.

Ritonavir induces many more significant shifts in comparison to the other protease inhibitors tested, Darunavir, Indinavir and Nelfinavir. Two of the hits CES2 and DHRS4 could be implicated in the metabolism of Ritonavir. The other stability-altered proteins may constitute a phenotype where lipid metabolism pathways are affected alongside Ca and H+ ion balance. There is support in the literature for effects on lipid metabolism and on the respiratory chain^26,27^. Proteins involved in Lipid metabolism being thermally shifted by Ritonavir include FABP1, MTTP, ACOX1, ACOX3, HADHA, DECR2, AIFM1 and AIFM2. The thiol modification protein QSOX2. Ion channels LETM1 and SLC38A10. Calcium and zinc binding proteins HPCAL1 and NUCB1, Synaptic vesicle membrane protein VAT-1 homolog (VAT1) and finally, Cytosolic 5’-nucleotidase 3A (NT5C3A) that dephosphorylates CMP and 7m-GMP. It should be noted however, that most of the proteins shifted by Ritonavir are also shifted, albeit to a lower extent among the other protease inhibitors as well as a resemblance to the nonnucleoside inhibitor Delavirdine. A possible explanation is that Ritonavir has a faster cellular uptake or induction of cellular phenotypic effects resulting in a significantly stronger shift in these patterns than other compounds.

Baloxavir marboxil, the antiviral medication for treatment of influenza A and B, has quite distinct proteome stability alteration pattern. Baloxavir marboxil protein hits do not overlap with those of the other compounds. The proteins shifting include Thrombospondin-1 (THBS1), an adhesive glycoprotein that mediates cell-to-cell and cell-to-matrix interactions. Pyruvate Dehydrogenase protein component (PDHX), Mitochondrial ribosomal protein L39 (MRPL39), Lon protease (LONP1) an ATP dependent serine protease, Trifunctional enzyme subunit alpha (HADHA) and long-chain-fatty-acid-CoA ligase 1 (ACSL1). In our experience, such effects to the proteome are indicative of an oxidative stress response.

The remaining compounds either induce no shifts or do so for very few proteins. The latter make up cluster 2 in Figure 2 where the lack of pronounced molecular fingerprint does not allow for further division into separate or unique groupings. Lamivudine treatment resulted in in a stabilizing shift for DCK, which is known to be responsible for the intracellular phosphorylation of the drug^28^ which provide confidence of cellular uptake. These data may well constitute useful information when taken in the context of further study.

## Discussion

This study intended to help better understand any off-target effects of Remdesivir and Chloroquine as two prominently repurposed drugs for targeting SARS-CoV-2, with the view to identify potential biological inroads for further investigation.

This is an intact cell study and therefore conducted in a highly biological context. In that, proteins exist at their endogenous expression and environment. The relative amounts of analytes, nucleotide and metabolites represent levels commensurate to healthy unmodified cultured cells. In the CETSA MS platform, we identify both stabilized and destabilized proteins after treatment with these drug molecules. A stabilizing shift is often attributed to a direct binding event. Similarly a destabilizing shift can also be caused by a direct binding event if the molecular interaction causes the target to be less thermodynamically favorable. Additionally, destabilizing events can be caused by the usurping of a native substrate or the removal of a complex of protein-protein interactions as a secondary downstream effect.

The host targets identified represent a wide variety of biological processes. It is important to note that these data are included in this study to offer an unrevised perspective at potential off-targets. A thorough understanding of the relevance to viral infection is a significant undertaking and well beyond the scope and timelines of this study. The term off-targets or unintended targets is employed here, meaning not the primary target. Although it can also be the case that an interaction with a host protein is essential for the efficacy of the drug, as is the case with Lamivudine and DCK. In this light, there are other targets involved in nucleotide regulation such as RNR, SAMHD1 and ADK in particular that we would have expected the cellular presence of GS-441524 and other nucleotide analogues to have affected. In contrast, these regulatory proteins were not identified as hits which is a surprising outcome given in our experience ADK is known to shift upon binding of substrate.

There are no previous studies using CETSA MS to comparatively analyze a panel of anti-viral compounds. Given the primary purpose of the majority of these drugs is to interact or inhibit viral proteins, there was no expectation that common host targets would be identified.

In contrast, the Chloroquine molecules are known to have substantial effects to the endosomal compartment and the expectation was significant and broad shifts in these samples that was not observed. Aside from the described shifts, the bulk of cluster 2 represented less defined changes to a broad range of biological activities, not allowing for a definitive molecular fingerprint to be elucidated.

This study was designed to identify previously unidentified proteins that could have critical importance for the reported activity of Remdesivir and other compounds in the context of COVID-19. The inclusion of a panel of molecules allows for the cross comparison against hits specific to one molecule, which has facilitated the novel finding that Remdesivir uniquely destabilizes Trip13.

The function of Trip13 does not lend itself to that of an obvious benefit or hinderance to viral infection, as would be considered by a protein with known host innate viral immunity activity. But the fact it interacts with nucleotides and forms a homohexamer which if diminished removes activity, lends it to the possibility the interaction with Remdesivir may in fact be tangible^29^.

Further in vitro biophysical investigation probing the interaction could elucidate evidence into the role of Trip13 in Remdesivir therapy. The functional relevance of such an interaction in the context of viral infected tissue could yield crucial information as to whether its potential off target behavior is tolerable, beneficial or indeed a hindrance to the molecule’s efficacy against Sar-CoV-2.

This study has highlighted the power of utilizing unbiased whole proteome approaches and the information that can be rapidly gained from describing proteome wide target engagement of drug molecules.

